# Differential *O*-Glc elongation on the specific EGF repeat within a ligand binding domain regulates NOTCH1 signaling

**DOI:** 10.1101/2025.02.20.639268

**Authors:** Yohei Tsukamoto, Kazuhiro Aoki, Wataru Saiki, Natsumi Tsukamoto, Koki Kato, Yohei Hosokawa, Rie Sato, Yuki Fujita, Kana Fukazawa, Yuuki Kurebayashi, Yusuke Urata, Sae Uchiyama, Weiwei Wang, Akira Minami, Tadanobu Takahashi, Michael Tiemeyer, Yoshiki Narimatsu, Tetsuya Okajima, Hideyuki Takeuchi

**Author notes:** **Corresponding author: *** Hideyuki Takeuchi, **Email:** (H.T.).

## Abstract

Three types of *O*-linked glycosylation - *O*-glucose, *O*-fucose, and *O*-*N*-acetylglucosamine - are crucial for the function of Notch receptors, which regulates critical cell fate determination processes in a wide variety of contexts. *O*-Glucose glycans are added to serine residues located between the first and second conserved cysteines within the epidermal growth factor (EGF) -like repeats in the Notch extracellular domain. Previously, *O*-glucose glycans were shown to be extended to a trisaccharide structure with two xyloses via an α1-3 linkage. Our recent studies, however, indicated that the *O*-glucose glycan on NOTCH1 EGF10 can be extended by hexose and Neu5Ac. Here, we demonstrated that this hexose- and Neu5Ac-extended glycan has a 3’-sialyllactose-like structure synthesized by specific members of two isoenzyme families, B4GALT1 and ST3GAL4. Using mass spectrometry, we identified this modification exclusively on NOTCH1 EGF10 and NOTCH3 EGF9 (equivalent to NOTCH1 EGF10), with no detection in any other EGF domains in NOTCH1-3. Sequence comparison and mutagenesis experiments identified one amino acid at position -2 of the fourth cysteine in the EGF domain as crucial for the galactose elongation of *O*-glucose glycans. We further demonstrated that this site-specific elongation of *O*-glucose on NOTCH1 EGF10 significantly impact ligand binding and signal transduction of NOTCH1. Our findings contribute to the understanding of the intricate regulatory mechanisms of Notch receptor function mediated by distinct positions and structures of *O*-glycans.

**Significance Statement:** Notch, a signaling receptor critical for development, differentiation, and homeostasis in multicellular organisms, undergoes multiple post-translational glycan modifications to regulate its function. While *O*-glucose glycan elongation by xylose was previously known, we identified an *O*-glucose glycan extended by galactose and Neu5Ac on specific epidermal growth factor (EGF) -like repeats of certain Notch paralogs, NOTCH1 and NOTCH3. As far as we know, this structure has not been reported on any other proteins. We identify the enzymes involved in the biosynthesis of this glycan and demonstrate that modifications at a specific site on Notch direct receptor functions, including ligand binding and Notch signaling.

## Introduction

Notch receptors are crucial for the development and homeostasis of multicellular organisms (3). Biochemical and genetic analyses have shown that *O*-glycosylation regulates Notch receptor function (4, 5). Abnormal *O*-glycosylation has been linked to several human diseases that are caused by mutations in glycosyltransferases involved in Notch glycosylation (6–10). Notch receptors are type I membrane protein with four paralogs in mammals, possessing 29 to 36 epidermal growth factor-like (EGF) repeats in their extracellular domain. These Notch receptors share five canonical ligands, three Delta-like ligands (DLL1, DLL3 and DLL4), and two Jagged ligands (JAG1 and JAG2) (11). Ligands on neighboring cells bind to Notch receptors, which unfolds the Notch negative regulatory region and exposes the cleavage site of ADAM10/17 metalloprotease (S2 cleavage). After S2 cleavage, the Notch transmembrane domain is cleaved by γ-secretase complex (S3 cleavage), releasing the intracellular domain into the cytoplasm (12). This domain then enters the nucleus, where it forms a complex with DNA-binding proteins such as RBPj/CSL to regulate the transcription of downstream genes such as Hes and Hey families (13).

EGF repeats in the Notch extracellular domain are modified by three distinct types of *O*-glycans. *O*-Glucose (Glc), *O*-fucose (Fuc), and *O*-*N*-acetylglucosamine (GlcNAc)-type glycans are added at a specific position within a consensus sequence. *O*-Glc is attached to a serine residue in the sequence C^1^-X-<u>S</u>-X-(P/A)-C^2^ by POGLUT1 (where X is any amino acid, and the modified residue is underlined) (5) or to a serine residue in C^3^-X-N-T-G-<u>S</u>-(F/Y)-X-C^4^ by POGLUT2 and POGLUT3 (14, 15). *O*-Fuc is added to a serine or threonine residue in the sequence C^2^-X-X-X-(<u>S/T</u>)-C^3^ by POFUT1(16), and *O*-GlcNAc is attached to a serine or threonine residue in the sequence C^5^-X-X-(G/S/T)-(Y/F/T)-(<u>S/T</u>)-G-X-X-X-C^6^ by EOGT (17, 18). These *O*-glycosylation sequence motifs are each found in specific EGF repeats of Notch and the other proteins. Each *O*-glycan can be elongated to different structures (19). All three types of *O*-glycosylation on mouse NOTCH1 have been analyzed by mass spectrometry. These modifications show site-specific variations in their occupancy and elongation. Notably, such variations are observed even for the same glycan species. (18, 20, 21). The importance of EGF-specific elongation of *O*-Fuc glycans for the function of Notch receptors is well demonstrated. In *Drosophila*, Fringe-mediated elongation of *O*-Fuc increases Notch signaling from delta-like ligands and reduces signaling from Serrate-like ligands (22). Similarly, in mammals, the elongation of *O*-Fuc by Fringe enhances NOTCH1 and NOTCH2 signaling derived from DLL1, and NOTCH2 signaling from DLL4, while reducing NOTCH1 and NOTCH2 signaling derived from JAG1 (21, 23). These findings highlight the importance of specific glycan structures at particular EGF domains in activation of Notch, although many aspects remain unresolved.

Xylose is added to *O*-Glc by GXYLT1 and GXYLT2 in an α1-3 linkage (24) and the Xyl-Glc disaccharide is further elongated by XXYLT1 to form the Xylα1-3Xylα1-3Glc trisaccharide (25). Xylosyl elongation has been shown to suppress Notch activation in *Drosophila* (26). We generated *GXYLT1/2*-double and *XXYLT1* single knockout (KO) HEK293T cells and analyzed their phenotypes. We showed that xylosyl elongation is required for cell surface Notch expression at high levels. Surprisingly, in wild-type HEK293T cells, we found that *O*-Glc in EGF10 of NOTCH1 was elongated with hexose and Neu5Ac instead of xylose (Fig. 1A) (20). As far as we know, the Neu5Ac-hexose-Glc trisaccharide, a structure also found in the glycolipid GM3, has not been reported in any glycoproteins.

**Fig. 1.**
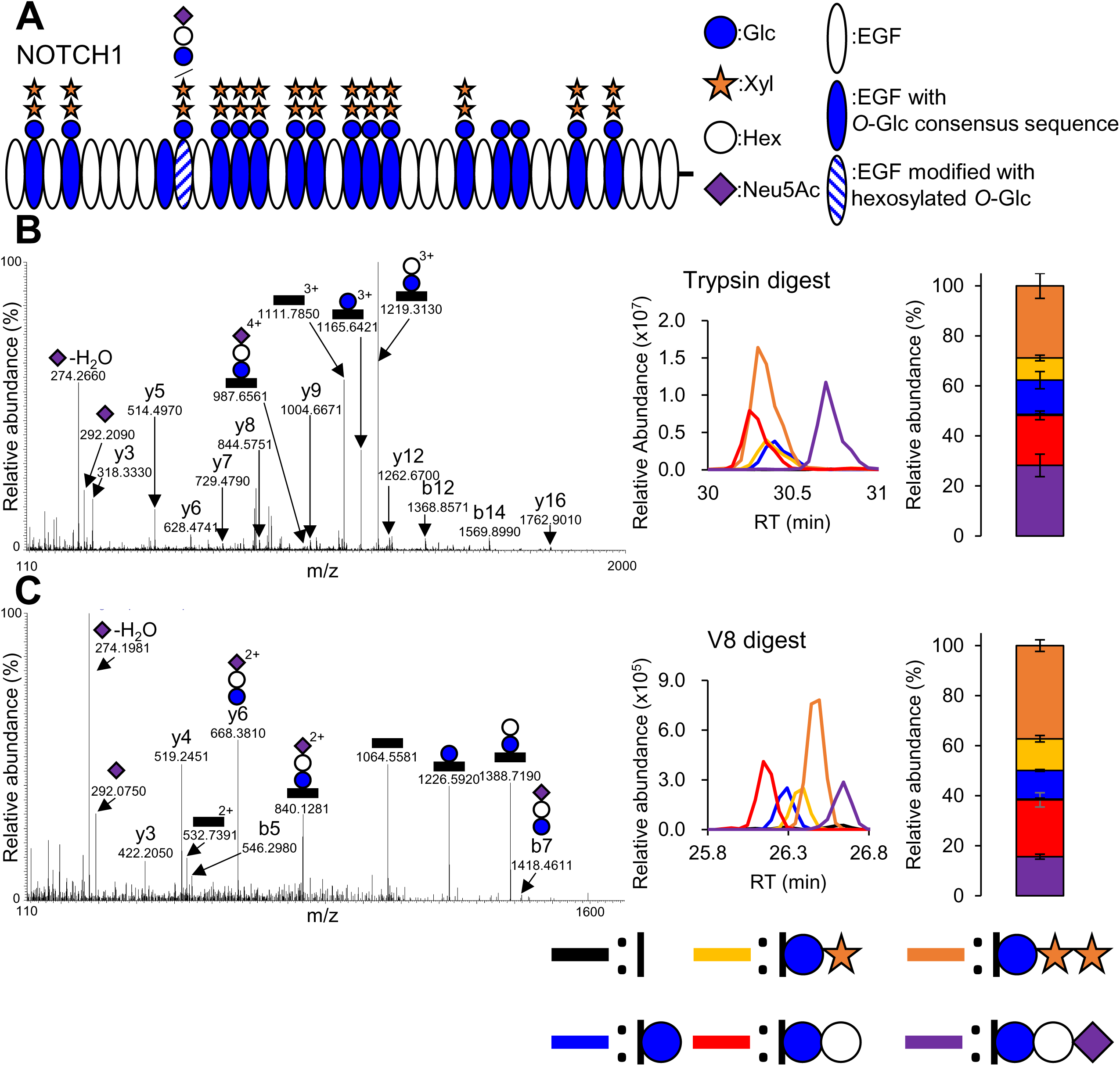
NOTCH1 EGF10 is modified by *O*-Glc elongated by galactose and Neu5Ac. Mass spectrometry data of protease-digested NOTCH1 EGF10. (A) Summary of *O*-glucosylation on NOTCH1 (20, 34, 62). Major glycoforms at each EGF repeat are shown. Blue circle: glucose; orange star: xylose; white circle: hexose; purple diamond: Neu5Ac; white oval: EGF; blue oval: EGF with *O*-Glc consensus sequence; blue and white stripe oval: EGF modified with hexosylated *O*-Glc. (B) Tryptic digest of mouse NOTCH1 EGF1-12. *Left*: MS/MS spectrum of peptide, 366-TGLLCHLNDACISNPCNEGSNCDTNPVNGK-395; *middle*: Extracted ion chromatograms (EICs) of each (glyco)peptide; *right*: Stoichiometry of each glycoform shown as a bar graph (N = 3). (C) V8 digest of mouse NOTCH1 EGF1-12. *Left*: MS/MS spectrum of peptide, 375-ACISNPCNE-383; *middle*: EICs of each (glyco)peptide; *right*: Stoichiometry of each glycoform shown as a bar graph (N = 3). In the left panels, representative peaks are annotated. The color coding of EICs and bar graphs is shown at the bottom. Black: non-*O*-glucosylated peptide; blue: peptide with Glc monosaccharide; yellow: peptide with Xyl-Glc disaccharide; orange: peptide with Xyl-Xyl-Glc trisaccharide; red: peptide with Hex-Glc disaccharide; purple: peptide with Neu5Ac-Hex-Glc trisaccharide. Error bars represent the standard error of the mean (SEM).

In this study, we investigated the biosynthesis and function of this previously uncharacterized glycan Neu5Ac-hexose-Glc on EGF10 of NOTCH1 within the ligand binding region (27, 28). By combining our mass spectrometry and the genetic glycoengineering strategy (29), we determined that this EGF-specific glycan structure is specifically synthesized by distinct members of glycosyltransferase isoenzyme families, B4GALT1 and ST3GAL4. We further demonstrate that this elongated sialylated *O*-glycan structure negatively regulates NOTCH1 activation by reducing its binding affinity for the DLL1 ligand.

## Results

### Hexosyl and sialyl extended *O*-Glc glycan modifies NOTCH1 EGF10

To investigate glycan modifications of NOTCH1, we expressed and purified mouse NOTCH1 EGF1-12 proteins from HEK293T cells, digested them with trypsin or V8 protease, and analyzed the digests by mass spectrometry. In trypsin digests, we found a peptide 366-TGLLCHLNDACISNPCNEGSNCDTNPVNGK-395 from EGF10 modified with *O*-Glc glycans elongated by hexose or by hexose and Neu5Ac in addition to *O*-Glc glycans elongated with one or two pentose residues (Fig. 1B). V8 protease digestion further refined the modification site, revealing a peptide 375-ACISNPCNE-383 with similar glycoforms (Fig. 1C). The presence of fragment ions y6 and b7 strongly suggested that an *O*-Glc glycan was attached to the serine residue at position 378. We compared the extracted ion chromatograms (EICs) of *O*-Glc-modified peptides derived from EGF10 of mouse and human NOTCH1. The analysis revealed similar levels of hexose- and xylose-extended forms between the two species (Fig. 1).

*O*-Glc glycans elongated by hexose were detected on EGF10 of NOTCH1 expressed in various human cell lines including oral squamous cell carcinoma cells (SAS), pancreatic adenocarcinoma cells (ASPC1), and leukemia cells (Jurkat) cells. These results suggest that hexosyl elongation of *O*-Glc glycans is a common phenomenon in humans, but not tissue-specific (Fig. S1). The disappearance of all *O*-Glc glycoforms upon substituting serine at position 378 with alanine demonstrated that both hexosyl and xylosyl elongation occur on *O*-Glc attached to S378 (Fig. S2A). Knocking out both *GXYLT1* and *GXYLT2* abolished xylsoyl elongation and did not affect hexosyl elongation. Overexpression of *GXYLT1/2* genes in GXYLT1/2-double knockout (DKO) cells increased xylosyl elongation and decreased hexosyl elongation of *O*-Glc glycans (Fig. S2B, C).

### *O*-Glc glycans can be extended to form lactose and 3’-sialyllactose structures

To investigate the structure of the hexosyl glycans on NOTCH1 EGF10, chemical analysis was conducted. Using the method from a previous report (30, 31), we analyzed *O*-Glc glycans extended by a hexose and Neu5Ac released from the tryptic glycopeptide, 366-TGLLCHLNDACISNPCNEGSNCDTNPVNGK-395, of NOTCH1 EGF6-10 by reductive β-elimination. The released, permethylated glycan and a reduced, permethylated 3’-sialyllactose standard were subjected to nanoelectrospray ionization-multidimensional mass spectrometry (NSI-MSn), yielding fragmentation patterns consistent with 3’-sialyllactose (Fig. 2A and Fig. S3) (30). Signature NSI-MSn fragment ions were detected at MS4 for the glycan released from NOTCH1 EGF6-10 and for the 3’-sialyllactose standard, including ^3,5A^Gal at m/z 109, ^1,3X^Gal at m/z 137, and ^2,3X^Gal at m/z 181 which are diagnostic for a C3-substituted hexose. Therefore, the terminal sialic acid is attached at the C3 position of the internal hexose residue (30). Furthermore, the glycopeptides from the EGF10 of NOTCH1 expressed in *GXYLT1/2* DKO cells were digested with glycosidases. Sialidase (cleaving all Siaα2-3/6/8/9 linkages) treatment markedly reduced the sialylated trisaccharide and further digestion with β4-galactosidase decreased the disaccharide to the hexose monosaccharide (Fig. 2B-D), strongly supporting the interpretation of the 3’-sialyllactose structure. Hereafter, the galactosyl elongation and xylosyl elongation processes are referred to as the 3’-sialyllactose pathway and xylose pathway, respectively.

**Fig. 2.**
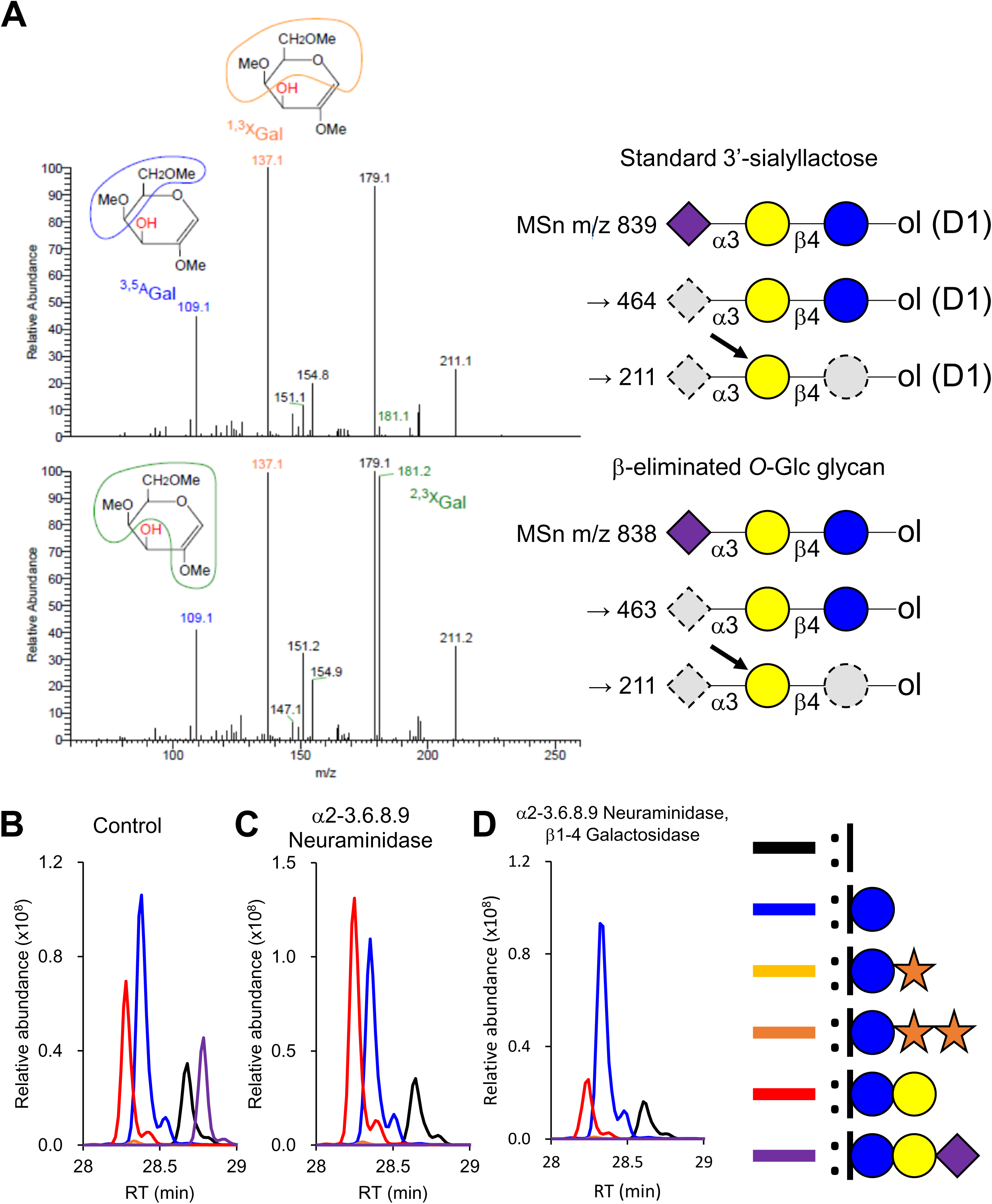
The *O*-Glc glycan is extended to form a 3’-sialyllactose-like structure. (A) The standard 3′-sialyllactose (Neu5Acα1-3Galβ1-4Glc) was permethylated following reductive β-elimination with sodium borodeuteride (NaBD_4_), resulting in a mass shift of +1 Da (m/z 839 in MS1). After β-elimination with sodium borohydride (NaBH_4_), the trisaccharide released from trypsin-digested glycopeptides, 366-TGLLCHLNDACIANPCNEGSNCDTNPVNGK-395, containing the EGF10 *O*-Glc site from NOTCH1 EGF6-10 was detected as its permethylated form (m/z 838 in MS1). For NSI-MSn analyses, permethylated trisaccharides were infused into a mass spectrometer in the presence of lithium ions, inducing linkage-specific sugar cross-ring cleavages (*right*), as previously reported. Arrows indicate sugars fragmented in MS4 analyses (*right*). The MS4 spectra (MS4 838@463@211) of the released trisaccharide correspond to those of the standard 3′-sialyllactose (Neu5Acα1-3Galβ1-4Glc) (*left*). Both spectra yields signature fragments such as ^3,5A^Gal at m/z 109 (blue), ^1,3X^Gal at m/z 137 (yellow), and ^2,3X^Gal at m/z 181 (green) identifying the presence of a 3-substituted internal hexose residue (58). (B-D) Glycoproteomics analysis of glycosidase-treated trypsin digest of mouse NOTCH1 EGF1-18. EICs of mouse NOTCH1 EGF10 glycopeptides with or without glycosidase treatment. B: Untreated control. C: α2-3.6.8.9-neuraminidase treatment. D: α2-3.6.8.9-neuraminidase treatment followed by β1-4-galactosidase treatment. The color coding of EICs is shown in the bottom right. Black: non-*O*-glucosylated peptide; blue: peptide with Glc monosaccharide; yellow: peptide with Xyl-Glc disaccharide; orange: peptide with Xyl-Xyl-Glc trisaccharide; red: peptide with Hex-Glc disaccharide; purple: peptide with Neu5Ac-Hex-Glc trisaccharide. Blue circle: glucose; orange star: xylose; yellow circle: galactose; purple diamond: Neu5Ac; Sugars that are cleaved off through neutral loss are shown as dashed gray symbols.

### *B4GALT1* and *ST3GAL4* are involved in the biosynthesis of the 3’-sialyllactose elongated *O*-Glc glycan

To identify the glycosyltransferase isoenzymes responsible for the 3’-sialyllactose pathway, we analyzed NOTCH1 proteins expressed in wild-type HEK293 cells and isogenic cells with various combinations of KO of *B4GALT*, *ST3GAL,* and/or *ST6GAL* isoenzyme genes by mass spectrometry (Fig. 3). Combinatorial KO of *B4GALT1-4* resulted in the loss of extended *O*-Glc glycans (di- and trisaccharide peaks) in the 3’-sialyllactose pathway, while DKO of *B4GALT5/6* did not. In *B4GALT1-4*-deficient cells, *O*-Glc glycans elongated by galactose were not detected, while they were in *B4GALT5/6* deficient cells as wild-type cells (Fig. 3A, B). Further KO of individual *B4GALT* genes demonstrated that KO of *B4GALT1* alone abrogated synthesis of extended *O*-Glc (Fig. 3C). Combinatorial KO of *ST3GAL1-6* with or without *ST6GAL1/2* abrogated Neu5Ac addition, whereas *ST6GAL1/2* DKO had no effect (Fig. 3D). Furthermore, only *ST3GAL4*-deficient cells showed a marked reduction in Neu5Ac addition (Fig. 3E). We confirmed these results by overexpression studies, demonstrating that reintroduction of cDNA constructs for B4GALT1 and ST3GAL4 rescued the *O*-Glc elongation by galactose and Neu5Ac, respectively (Fig. S4).

**Fig. 3.**
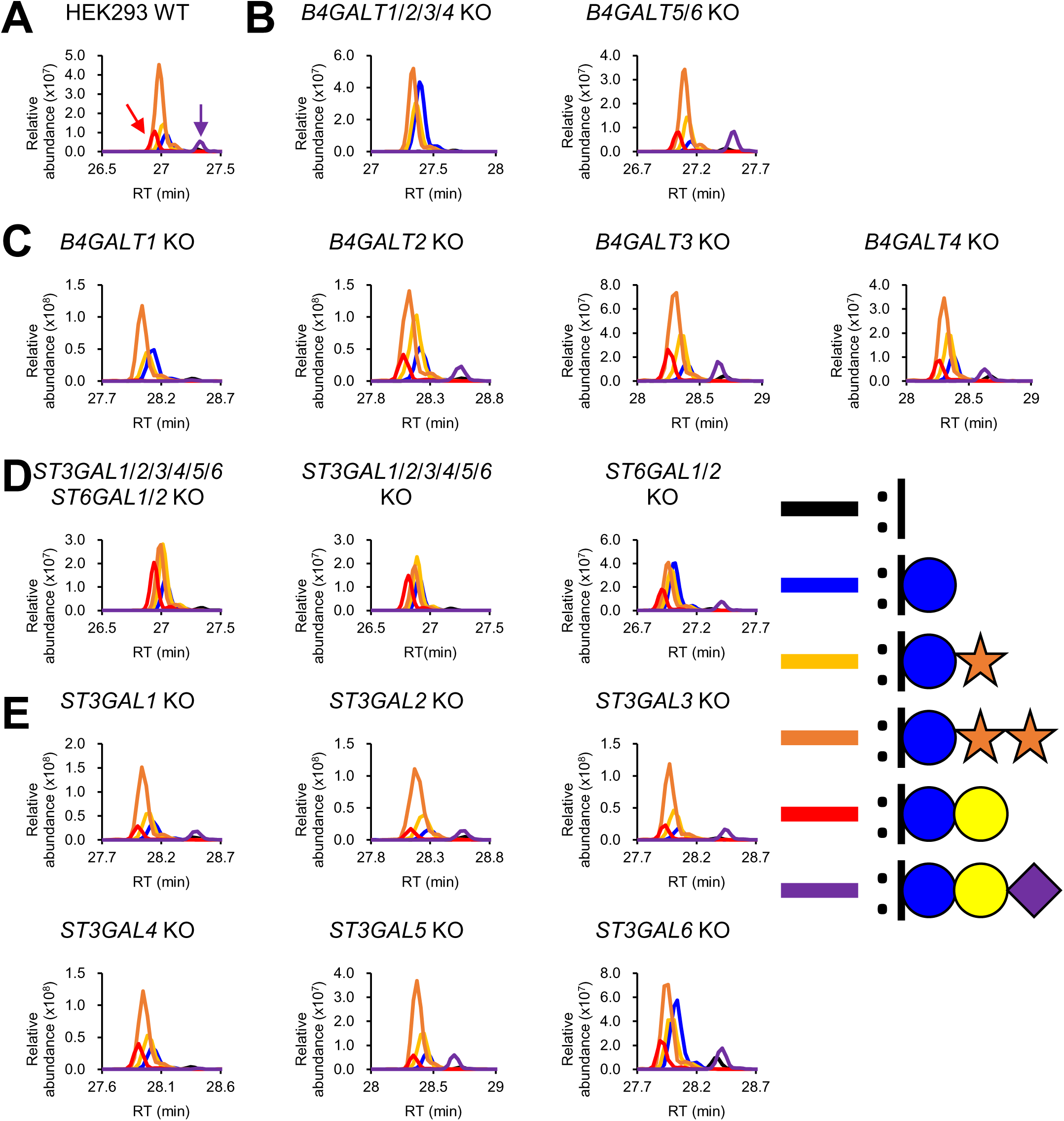
B4GALT1 and ST3GAL4 are involved in the *O*-Glc glycan biosynthesis. Mouse NOTCH1 EGF6-10 was digested by trypsin and analyzed by mass spectrometry. For the analysis with *B4GALT5/6* KO cells, mouse NOTCH1 EGF1-12 was digested by trypsin. EICs of (glyco)peptides, 366-TGLLCHLNDACISNPCNEGSNCDTNPVNGK-395, are shown. (A) Wild-type (WT) HEK293 cells. The red arrow indicates galactose-extended *O*-Glc glycan and the purple arrow indicates galactose- and Neu5Ac-extended *O*-Glc glycan. (B) Multiple *B4GALT* gene-deficient HEK293 cells. (C) Single *B4GALT* gene-deficient HEK293 cells. (D) Multiple *ST3GAL* and/or *ST6GAL* gene-deficient cells. (E) Single *ST3GAL* gene-deficient cells. The color coding of EICs is shown in the bottom right. Black: non-*O*-glucosylated peptide; blue: peptide with Glc monosaccharide; yellow: peptide with Xyl-Glc disaccharide; orange: peptide with Xyl-Xyl-Glc trisaccharide; red: peptide with Gal-Glc disaccharide; purple: peptide with Neu5Ac-Gal-Glc trisaccharide.

### The 3’-sialyllactose elongation pathway of *O*-Glc glycans is directed by the sequence context of NOTCH EGF repeats

The *O*-Glc glycans in the 3’-sialyllactose pathway were detected on mouse NOTCH1 EGF10 in wild-type HEK293T cells. Previous reports showed that these glycans were absent on most EGF repeats from mouse NOTCH1 and NOTCH2, even in *GXYLT1/2*-deficient cells although some EGF repeats, such as NOTCH1 EGF12, NOTCH2 EGF12, and NOTCH2 EGF33, were modified with those *O*-Glc glycans at varying levels (20), indicating EGF-specific glycosylation. EGF repeats among Notch paralogs are highly conserved, exemplified by the sequence of EGF10 (Fig. 4A) (32). We analyzed *O*-Glc glycosylation on mouse NOTCH2 and NOTCH3 by mass spectrometry. NOTCH2 EGF10 lacked *O*-Glc glycans in the 3’-sialyllactose pathway, whereas NOTCH3 EGF9, equivalent to NOTCH1 EGF10, was modified (Fig. 4B, C). Mass spectrometry analysis of human NOTCH1 EGF8-13 expressed in HEK293T cells revealed *O*-Glc glycans in the 3’-sialyllactose pathway on EGF10 (Fig. 4D). Reanalysis of published human NOTCH3 mass spectrometry data (33) confirmed similar modifications on EGF9 (Fig. 4E).

**Fig. 4.**
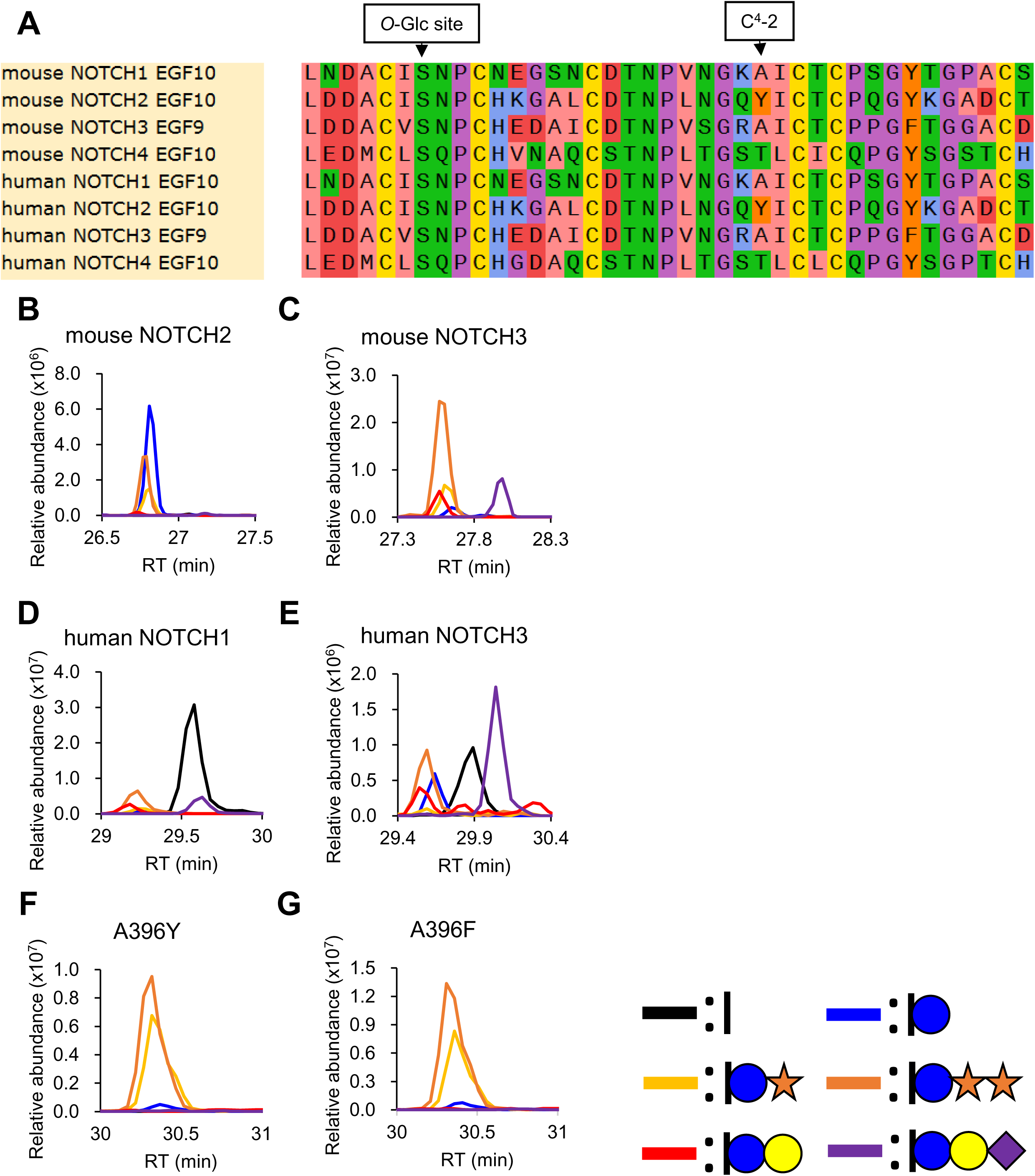
The *O*-Glc elongation in the 3’-sialyllactose pathway is specific to NOTCH1 EGF10 and NOTCH3 EGF9. (A) Sequence alignment of mouse and human NOTCH1 EGF10, NOTCH2 EGF10, NOTCH3 EGF9, and NOTCH4 EGF10 generated by snap gene software. The color coding of amino acids is as follows. Pink: hydrophobic; red: negatively charged; yellow: cysteine; green: hydrophilic; blue: positively charged; purple: conformationally special; orange: aromatic. (B-E) EICs of Notch protein tryptic digests. B: mouse NOTCH2 EGF10. C: mouse NOTCH3 EGF9. D: human NOTCH1. E: human NOTCH3 EGF9 (See also mouse NOTCH1 data shown in Fig. 1B). (F, G) Glycan analysis of A396Y (F)- and A396F (G)-mutant mouse NOTCH1 EGF1-12. The color coding of EICs is shown in the bottom right. Black: non-*O*-glucosylated peptide; blue: peptide with Glc monosaccharide; yellow: peptide with Xyl-Glc disaccharide; orange: peptide with Xyl-Xyl-Glc trisaccharide; red: peptide with Gal-Glc disaccharide; purple: peptide with Neu5Ac-Gal-Glc trisaccharide.

Alignment of EGF10 (and its equivalent) sequences from Notch paralogs indicates a difference in the amino acid located two residues before the fourth cysteine (C^4^-2 position) (Fig. 4A). In EGF repeats containing the O-Glc consensus sequence, aromatic residues at C^4^-2 are highly conserved (34). To investigate the effect of C^4^-2 amino acids on the galactosyl elongation, we generated A396Y/F mutants (mutations at C^4^-2 of EGF10) of mouse NOTCH1 EGF1-12 and expressed them in HEK293T cells. Mass spectrometry analysis showed that both mutants lacked galactosyl elongation on *O*-Glc at EGF10 (Fig. 4F, G). Additionally, Y85A and Y162A mutants (mutations at C^4^-2 positions in EGF2 and EGF4) of mouse NOTCH1 EGF1-12 reduced *O*-Glc glycosylation and xylosyl elongation, with only trace galactosyl elongation detected (Fig. S5). These results suggest that C^4^-2 amino acids are crucial for determining two distinct *O*-Glc glycan elongation pathways.

### The elongation of *O*-Glc glycans on NOTCH1 EGF10 regulates ligand binding and signaling

To investigate the impact of *O*-Glc glycan modifications on Notch function, we performed ligand binding assays with truncated mouse NOTCH1 EGF8-12 proteins containing ligand-binding sites. We prepared truncated mouse NOTCH1 with different *O*-Glc glycosylation patterns using three conditions: (1) *B4GALT1* KO HEK293 cells with POGLUT1 and GXYLT1 overexpression, (2) *GXYLT1/2* DKO HEK293T cells with POGLUT1 and B4GALT1 overexpression, and (3) wild-type HEK293T cells with POGLUT1 overexpression (Fig. S6). The *O*-Glc glycosylation of the EGF10 glycopeptides was confirmed by mass spectrometry (Fig. S6). Then the purified mouse NOTCH1 proteins were incubated with HEK293T cells transiently overexpressing the canonical Notch ligands DLL1, DLL4, or JAG1, and ligand-binding was measured by flow cytometry. Compared to wild-type proteins, those lacking galactose elongation exhibited higher binding to all of the ligands, while those lacking xylose elongation were unchanged or slightly reduced (Fig. 5 and Fig. S6). We also measured binding of soluble DLL1, DLL4, and JAG1 to HEK293T cells transiently expressing wild-type or A396Y/F mutants of full-length NOTCH1. The wild-type and both mutants were expressed on the cell surface at similar levels (Fig. S7), and A396Y/F mutant-expressing cells showed markedly increased binding to DLL1, but not to DLL4 and JAG1 (Fig. S8).

**Fig. 5.**
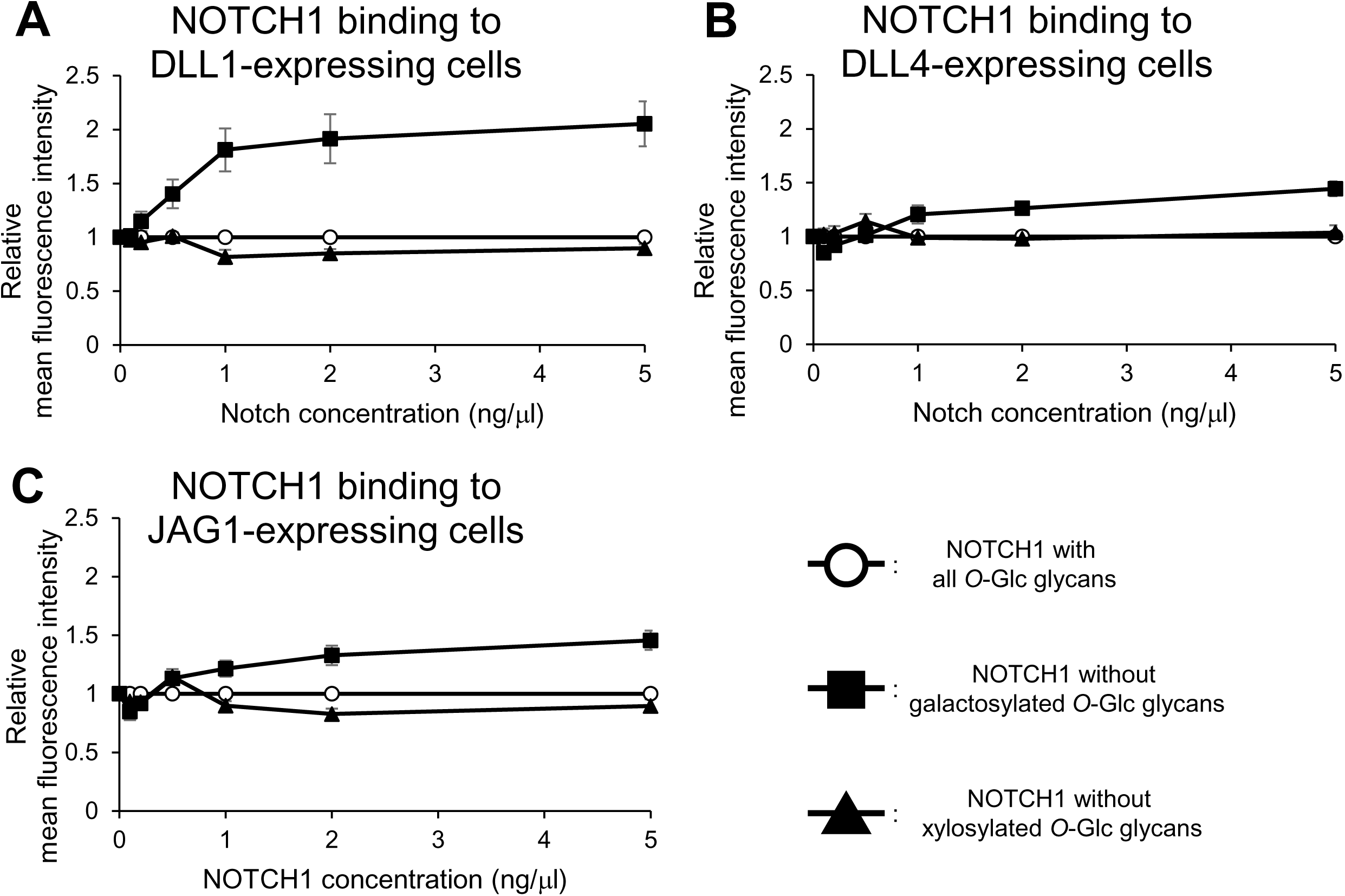
Differential *O*-Glc elongation on NOTCH1 EGF10 plays an important role in NOTCH1-ligand binding regulation. (A-C) HEK293T cells overexpressing full-length Notch ligands DLL1 (A), DLL4 (B), or JAG1 (C) were incubated with mouse NOTCH1 EGF8-12 (0-5 ng/μL). Circles: mouse NOTCH1 EGF8-12, produced in *POGLUT1*-overexpressing wild-type HEK293T cells, containing both galactose- and xylose-elongated *O*-Glc glycans; squares: mouse NOTCH1 EGF8-12, produced in *POGLUT1*- and *GXYLT1*-overexpressing *B4GALT1* KO cells, lacking galactose-elongated *O*-Glc glycans; triangles: mouse NOTCH1 EGF8-12, produced in *POGLUT1*- and *B4GALT1*-overexpressing *GXLYT1/2* KO cells, lacking xylose-elongated *O*-Glc glycans. All experiments were performed in triplicate and relative mean fluorescence intensity is shown on the y-axis. The intensity of mouse NOTCH1 EGF8-12 containing both galactose- and xylose-elongated *O*-Glc glycans was set as 1.0. The glycan composition of each NOTCH1 protein is shown in Fig. S6.

Since Notch receptor signaling is normally triggered by ligand binding, we hypothesized that changes in *O*-Glc glycan modifications on EGF10 could also affect the signaling. B4GALT1 and ST3GAL4 synthesize elongated *O*-Fuc and/or *O*-GlcNAc glycans on EGF repeats including EGF10 in NOTCH1 (35–37). To examine the role of EGF10-specific *O*-Glc glycan in the 3’-sialyllactose pathway, we performed signaling assays using HEK293T cells overexpressing wild-type or A396Y/F mutant of full-length NOTCH1. NOTCH1-expressing cells were stimulated by ligands coated on plates. *HES5* gene expression analysis by RT-qPCR and luciferase reporter assays were used as signaling readouts for NOTCH1 activation. The results showed significantly higher *HES5* gene expression and luciferase reporter activity in A396Y/F mutant-expressing cells stimulated by DLL1 compared to wild-type NOTCH1. However, there were no significant changes observed with DLL4 or JAG1 stimulation (Fig. 6A-E).

**Fig. 6.**
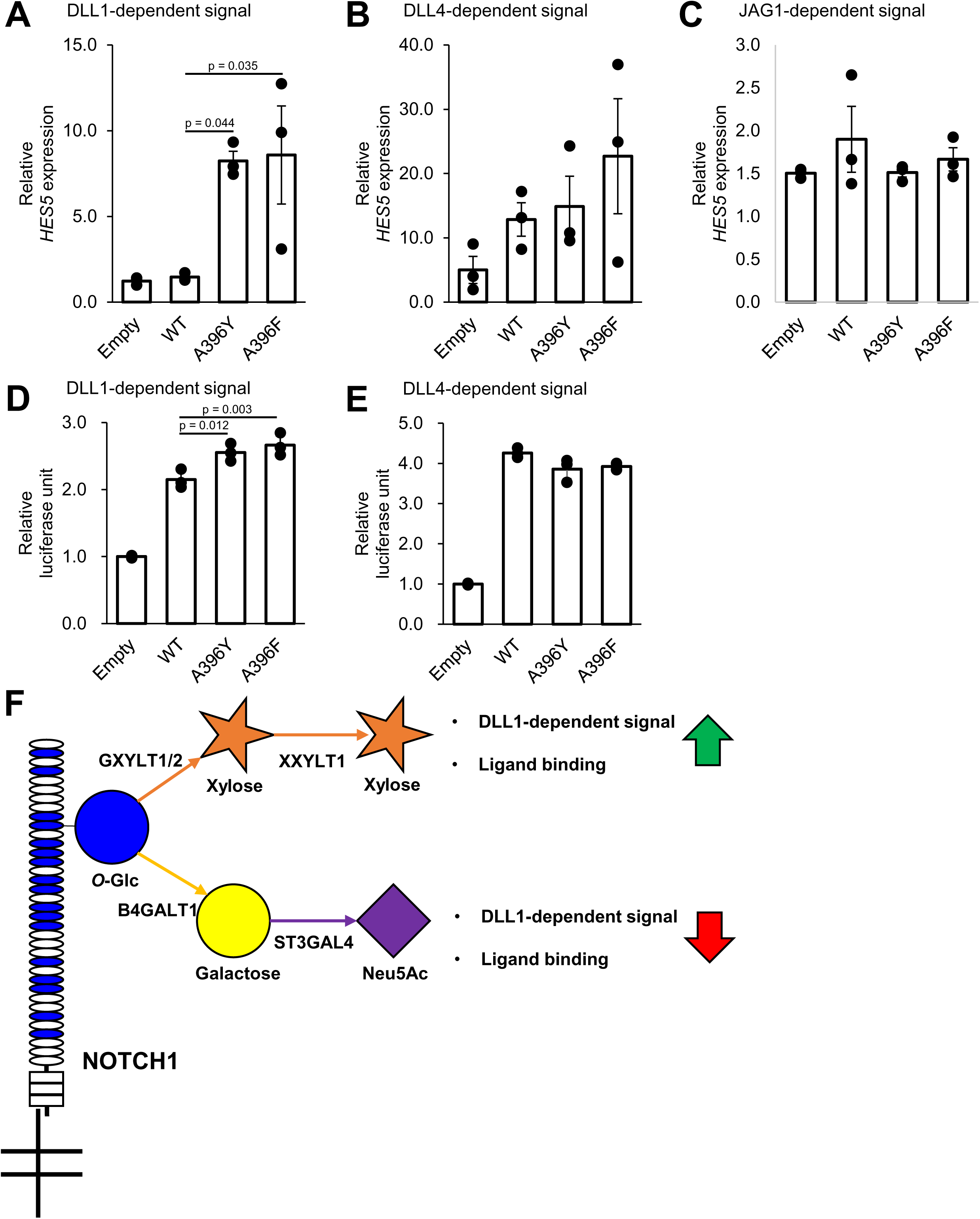
Differential *O*-Glc elongation on NOTCH1 EGF10 regulates NOTCH1 signaling. (A-C) RT-qPCR analysis of *HES5* gene expression. Empty vector, wild-type (WT), or mutant full-length NOTCH1-overexpressing cells were seeded on ligand-coated plates. A: DLL1-Fc stimulation (N = 3). B: DLL4-Fc stimulation (N = 3). C: JAG1-Fc stimulation (N = 3). The Ct value of *HES5* was normalized by the GAPDH Ct value. Negative control IgG-Fc-stimulated cells were set as 1.0. Statistical analysis was performed using one-way ANOVA followed by Tukey’s test. Error bars represent SEM. (D, E) Luciferase reporter assay. Empty vector, WT, or mutant full-length NOTCH1-overexpressing cells were seeded on ligand-coated plates. D: DLL1 Myc-His stimulation (N = 3). E: DLL4 Myc-His stimulation (N = 3). The relative luciferase unit of negative control IgG-Fc-stimulated cells was set as 1.0. Statistical analysis was performed using one-way ANOVA followed by Tukey’s test. Error bars represent SEM. (F) Summary of this study. The schematic of NOTCH1 and *O*-Glc glycans is shown. Oval: EGF repeats; blue oval: EGF repeats with *O*-Glc consensus sequence; blue circle: glucose; yellow circle: galactose; orange star: xylose; purple diamond: Neu5Ac; green arrow: upregulation; red arrow: down-regulation.

## Discussion

Our study discovered a unique 3’-sialyllactose *O*-Glc elongation pathway targeting highly specific Notch EGF repeats. We used genetic engineering in HEK293 cells to demonstrate that the elongations of *O*-Glc by galactose and subsequently Neu5Ac are regulated by distinct glycosyltransferase isoenzymes, B4GALT1 and ST3GAL4, out of these large isoenzyme families. We demonstrate that the *O*-Glc 3’-sialyllactose elongation occurs on highly specific EGF repeats in NOTCH1 and NOTCH3, which is regulated by unique sequence context in these repeats. Finally, we show that the 3’-sialyllactose elongation in EGF10 of NOTCH1 directs DLL1 binding and receptor activation.

EGF-specific *O*-Fuc and *O*-GlcNAc have been shown to carry sialylLac(NAc) (17, 38), and here we demonstrate that *O*-Glc in the 3’-sialyllactose pathway can also be modified with this structure. *O*-Glc 3’-sialyllactose elongation has never been found on any proteins as far as we know. Sialyllactose is a well-known glycan structure, primarily in the form of the ganglioside GM3 in glycolipids. The enzymes responsible for GM3 biosynthesis, B4GALT5, B4GALT6 (39), and ST3GAL5 (40), have been identified. Sialyllactose-like structures are found on many glycoproteins and are synthesized through common biosynthetic pathways. These pathways are regulated by isoenzyme families, where individual enzymes may exhibit varying degrees of specificity for certain structures. On the other hand, on NOTCH1, the main biosynthetic enzymes for sialyllactose are B4GATL1 and ST3GAL4, indicating that it is synthesized differently from glycolipids, in a manner similar to milk oligosaccharides (41) and *O*-GlcNAc glycans (35).

B4GALT1 typically adds galactose to GlcNAc via a β1-4 linkage (42), but with lactalbumin, it can add galactose to glucose via β1-4 linkage (43). The B4GALT1-lactalbumin complex structure shows that lactalbumin binding alters the conformation of B4GALT1, allowing it to recognize glucose instead of GlcNAc. Notably, lactalbumin is not expressed in HEK293T cells, suggesting that B4GALT1 adds galactose to *O*-Glc on specific EGF domains, such as NOTCH1 EGF10 and NOTCH3 EGF9, independently of lactalbumin. The overall structure of Notch is unclear, but unstable regions, such as EGF10, may act similarly to lactalbumin (44).

The enzymes responsible for xylosyl elongation of *O*-Glc glycans, GXYLT1/2 and XXYLT1, are thought to be localized to the endoplasmic reticulum (ER) (24, 25), whereas B4GALT1 and ST3GAL4 are Golgi-localized enzymes (45–47), suggesting that galactose and Neu5Ac elongation of *O*-Glc occurs in the Golgi apparatus. Thus, for galactose addition to *O*-Glc, xylosyl elongation should be suppressed in the ER. In A396Y/F mutants, almost all *O*-Glc glycans are xylosyl elongated, indicating that the alanine at C^4^-2 position of EGF10 inhibits xylosyl elongation, allowing the monosaccharide to reach the Golgi apparatus. Galactosyl elongation of *O*-Glc was observed in NOTCH1 EGF12, NOTCH2 EGF12, and NOTCH2 EGF33 in *GXYLT1/2*-deficient cells, though other *O*-Glc sites remain unaffected (20), indicating additional regulatory factors.

Ligand-binding assays show that the galactosyl elongation of *O*-Glc on EGF10 suppresses the interaction between NOTCH1 and DLL1. This suggests that the galactose-to-xylose ratio on *O*-Glc glycans finely regulates ligand binding. Previous analyses during the discovery of POGLUT1/rumi, experiments of *Poglut1* knockdown in mouse myoblast C2C12 cells did not suggest that *O*-Glc glycosylation affects ligand binding (5, 48). The apparent discrepancy between our current findings and those of previous studies may be attributed to differences in experimental systems. The former study was conducted in *Drosophila*, where *O*-Glc elongation modifications might differ fundamentally from those in mammalian systems. The latter study, on the other hand, investigated ligand binding to cells after knockdown of *Poglut1* in C2C12 cells with multiple Notch paralogs expressed, and it is possible that ligand binding specific to NOTCH1 was not detected. Structural analysis of the ligand-NOTCH1 complex revealed that serine 378, the *O*-Glc glycosylation site in EGF10, is opposite the ligand-binding interface (28), making the current findings unexpected. The most likely explanation is that *O*-Glc glycans in EGF10 affect the structural stability of the ligand-binding region. α-Linked xylose residues on *O*-Glc stabilize the EGF structure through intramolecular interactions involving calcium coordination (34), while galactosyl elongation, a β-linked sugar, may not provide similar stabilization. Since EGF10 lacks a calcium-binding motif, it is more sensitive to the conformational effects of sugar modifications, which likely influence ligand binding. Further structural and biochemical studies are needed for clarification.

Finally, this study identifies B4GALT1 and ST3GAL4 as the enzymes responsible for *O*-Glc in the 3’-sialyllactose pathway on NOTCH1 and NOTCH3 receptors. We also demonstrate that *O*-Glc in the 3’-sialyllactose pathway negatively regulates DLL1-dependent Notch signaling. The responsible enzymes are shared with elongation of *O*-Fuc and *O*-GlcNAc glycans on Notch (35, 49). Galactose elongation occurs after GlcNAc transfer by the fringe family enzymes to *O*-Fuc (37), and in CHO cells, this elongation regulates DLL1-dependent Notch activation (49). Future studies should explore whether elongation of *O*-Glc, *O*-Fuc, and *O*-GlcNAc are linked in regulating Notch activation to improve our understanding of glycosylation-based regulation. Nonetheless, this research underscores that variations in *O*-Glc glycans at a specific site of Notch receptors likely play a role in the fine regulation of receptor function.

## Materials and Methods

### Cell culture and transfection

Wild-type and previously reported isogenic glycosyltransferase gene-deficient HEK293 (2, 50) or HEK293T cells (20), were cultured in High Glucose Dulbecco’s modified Eagle Medium supplemented with 10% or 5% Fetal Bovine Serum, 100 U/ml penicillin, and 100 mg/ml streptomycin as previously reported (35). HEK293 or HEK293T cells were transfected with expression vectors using PEI MAX (Polyscience) as previously reported (20).

### Primers

All the primers used for subcloning, mutagenesis, and RT-qPCR in this study are listed in supporting information table S1.

### Plasmids

The mouse NOTCH1 EGF8-12 (amino acids 295-488, Uniprot Q01705) and human NOTCH1 EGF8-13 (amino acids 295-526, Uniprot P46531) fused with C-terminal myc and His tag were generated by PCR using KOD Fx Neo polymerase (TOYOBO) with pSectag2C plasmids encoding mouse NOTCH1 EGF1-36 and pcDNA3 plasmid encoding full-length human NOTCH1 as template and subcloned into expression vector pSectag2C using Hind III and Xho I cloning sites. The expression vector pcDNA4/TO_A_-CMV-mouse DLL1-Flag, pcDNA4/TO_A_-CMV-mouse JAG1-Flag were generated as follows. pTracer-CMV-mouse JAG1-Flag and pTracer-CMV-mouse DLL1-Flag (23) were digested with EcoR I and Xba I and the inserts were transferred to pcDNA4/TO_A_ (Invitrogen). The plasmid encoding C-terminally Myc-His-tagged mouse DLL4 was generated by PCR amplification using pCMV-sport6-mouse DLL4 (51) as a template and subcloning the product between EcoR I and Xba I sites in pcDNA4/TO_A_ (Invitrogen), in frame with the C-terminal Myc-His. The plasmid encoding mouse DLL1 ECD (∼Ser538) was created by amplifying the indicated region using pTracer-CMV-mouse DLL1-Flag as template and subcloning the product between EcoR I and Xho I sites in pcDNA4/TO_A_ (Invitrogen), in frame with the C-terminal Myc-His. The plasmid encoding mouse DLL4 ECD (∼Pro525) was created by amplifying the indicated coding region using pCMV-sport6-mouse DLL4 as template and subcloning the product between EcoR I and Xba I sites in pcDNA4/TO_A_ (Invitrogen), in frame with the C-terminal Myc-His. Successful cloning of all plasmids was confirmed by direct DNA sequencing. pTracer-CMV-mouse JAG1-Flag and pTracer-CMV-mouse DLL1-Flag were kindly sent from Dr. Robert S. Haltiwanger at University of Georgia.

Following plasmids were used as previously reported: pSectag2C mouse NOTCH1 EGF6-10 (52), pSectag2C mouse NOTCH1 EGF1-12 (35), pSectag2C mouse NOTCH1 EGF1-18 (52), pSectg2C mouse NOTCH1 EGF19-36 (52), pcDNA1 full-length mouse NOTCH1 Myc (21), pSectag2C mouse NOTCH2 EGF1-36 (14), pSectag2C mouse NOTCH3 EGF1-11 (35), pcDNA4 human POGLUT1 (34), pcDNA3 FLAG human GXYLT1-HA (53), pcDNA3 FLAG human GXYLT2-HA, pcDNA3 FLAG human XXYLT1-HA (25), pGEn2-DEST His-Avi-Super folder GFP-B4GALT1, pGEn2-DEST His-Avi-Super folder GFP-ST3GAL4 (35, 54), pMax-EGFP (20), human IgG-Fc-pRK5 (55). TP1-Luc, and gWiz-β-Gal (56) were kindly sent from Dr. Robert S. Haltiwanger at University of Georgia.

### Site-directed mutagenesis of mouse NOTCH1 protein expression vector

Mutations were introduced into pcDNA1 full-length mouse NOTCH1 Myc and pSectag2C mouse NOTCH1 EGF1-12 at the EGF2 C^4^-2 position (Y85A), EGF4 C^4^-2 position (Y162A), and EGF10 *O*-glucosylation site (S378A) and C^4^-2 position (A396Y and A396F) by site-directed mutagenesis PCR using KOD FX Neo polymerase (TOYOBO). The introduction of mutations was confirmed by DNA sequencing.

### Transient expression and protein purification of recombinant secreted forms of Notch proteins

HEK293 or HEK293T cells in 150 mm dishes were transiently transfected with 15 μg plasmid DNA and 75 μg polyethylenimine max (MW 40,000, Polysciences). After 4 hours of transfection, the medium was exchanged to OPTI-MEM (Gibco). After 72 hours of incubation, the culture supernatant was collected. To the collected culture media, NaCl and imidazole were added to a final concentration of 500 mM and 10 mM, respectively. The sample was applied to nickel-nitrilotriacetic acid (Ni-NTA) affinity column (Thermo Fisher Scientific), washed with 10□mM Tris-HCl pH 7.4 containing 650□mM NaCl and 10□mM imidazole, and eluted with TBS (10 mM Tris-HCl pH 7.4, 150 mM NaCl) containing 250 mM imidazole. Proteins used for ligand binding assay or signal assay were further dialyzed with TBS to remove imidazole.

### Mass spectrometry analysis

Sample preparation for mass spectrometry was performed as previously described (20, 35). Briefly, proteins were protease digested in solution or in gel by Trypsin Gold Mass Spectrometry Grade (V5280; Promega) or Endoproteinase Glu-C (V8, P8100S; NEB). Settings of Orbitrap Fusion Tribrid (Thermo Fisher Scientific) LC-MS were followed as described previously (35).

Mass spectral data were analyzed using Byonic software (Protein Metrics) and Xcalibur (Thermo Fisher Scientific) as previously reported with slight modification (35). For Byonic search, the following parameters were used. Trypsin or V8 digestion with a maximum of two possible missed cleavages. Cleavage sites were defined as C-terminal of Lys and Arg for Trypsin digestion, and C-terminal of Gln and Asp for V8 digestion. Carbamidomethylation on Cys as a fixed modification, and Oxidation on Met, Asn, and Asp, and Ammonia-loss on N-terminal Cys were set as common modification 1 (oxidation of Asp and Asn represents β-hydroxylation). The following glycan modifications were set as rare modification 1, Hex(1), Hex(1)Pent(1), Hex(1)Pent(2), Hex(2), Hex(2)NeuAc(1), Fuc(1), HexNAc(1), HexNAc(1)Hex(1), and HexNAc(1)Hex(1)NeuAc(1). For each peptide, a maximum of 4 common modifications and 2 rare modifications were considered. The MS/MS data analysis, scoring, and manual inspection of spectra were conducted as previously described (20, 35).

### Chemical analysis of β-eliminated galactosylated *O*-Glc glycan

Mouse NOTCH1 EGF6-10, containing the *O*-Glc consensus sequence only in EGF10, was expressed in *GXYLT1/2*-deficient HEK293T cells. Ni-NTA purified protein was digested by trypsin. Digested glycopeptides were separated by HPLC as previously described (57). Peaks containing galactose and Neu5Ac extended *O*-Glc glycan modified NOTCH1 EGF6-10 were collected by fraction collector. *O*-Glc glycans were eliminated from the peptide and permethylated as previously described (31). Permethylated glycans were analyzed by NSI-MSn as previously described (30, 58).

### Glycosidase treatment

β1-4 galactosidase treatment was performed as previously described (35). α2-3.6.8.9 Neuraminidase was obtained from Sigma (Proteomics Grade from Arthrobacter ureafaciens). The tryptic digest was incubated with α2-3.6.8.9 Neuraminidase in 50 mM sodium phosphate pH 6.0. After incubation, the reaction was terminated by heating at 100℃ for 5 min.

### Immunoblot analysis

pSectag2C mouse NOTCH1 EGF1-12 and human IgG-Fc for transient overexpression. IgG-Fc was used as secretion control. Protein expression and immunoblot was performed as previously described (20). iBright FL1500 (Thermo Fisher Scientific) was used for chemiluminescent signal detection. The antibodies used in this study were mouse anti-Myc antibody (1:2,000) (9E10; DSHB), goat anti-human IgG-Fc antibody HRP conjugated (1:50,000) (A80-104P; BETYHL) and anti-mouse IgG, HRP-linked Antibody (1: 10,000) (#7076P2; CST).

### Flow Cytometry

The cell surface expression levels of transiently overexpressed NOTCH1 were analyzed by flow cytometry using a FACS CANTOII Flow Cytometer, following previously reported methods (20). HEK293T cells, transiently overexpressing pcDNA1 full-length mouse NOTCH1 Myc or its mutants along with pMax-EGFP, were detached from the culture dishes using 1% BSA in PBS and washed with flow cytometry buffer (Hanks’ balanced salt solution containing 1% BSA, 1 mM CaCl_2_, and 0.02% NaN_3_). The cells were incubated on ice for 1 hour with 2 ng/µl of BV-conjugated anti-mouse NOTCH1 antibody (BioLegend, 130615), followed by two washes with 500 µl of flow cytometry buffer. Only singlet cells, filtered through a cell filter mesh, were analyzed using the FACS CANTOII. Live cells were gated based on Forward Scatter (FSC) and Side Scatter (SSC) properties, and the BV fluorescence intensity of the GFP-positive cell population was measured.

### Cell-Based NOTCH Binding Assay

The binding between soluble NOTCH and full-length ligands was analyzed using a FACS CANTOII Flow Cytometer as previously reported (59). HEK293T cells transiently overexpressing mouse DLL1, DLL4 or JAG1 along with pMax-EGFP were detached from the culture dishes using 1% BSA in PBS. The cells were washed with flow cytometry buffer (Hanks’ balanced salt solution containing 1% BSA, 1mM CaCl_2_, and 0.02% NaN_3_). Mouse NOTCH1 EGF8-12 Myc-His was preclustered with APC-conjugated anti-human c-Myc antibody (BioLegend, IC3696A) (1: 100) on ice for 1 hour and then added to cells. After incubating on ice for 1 hour, the cells were washed twice with 500 µl of flow cytometry buffer. Only singlet cells, filtered through a cell filter mesh, were analyzed using the FACS CANTOII. Live cells were gated based on FSC and SSC properties, and the APC fluorescence intensity of the GFP-positive cell population was measured.

### Cell-Based Ligand Binding Assay

The binding between full-length NOTCH and soluble ligands was analyzed using a FACS CANTOII Flow Cytometer as previously reported (49). HEK293T cells transiently overexpressing mouse NOTCH1 FL or its mutants along with pMax-EGFP were detached from the culture dishes using 1% BSA in PBS. The cells were washed with flow cytometry buffer. Rat DLL1-Fc, human DLL4-Fc, or human JAG1-Fc was preclustered with APC-conjugated anti-human IgG Fc antibody (BioLegend, 410712) (1: 100) on ice for 1 hour and then added to cells. After incubating on ice for 1 hour, the cells were washed twice with 500 µl of flow cytometry buffer. Only singlet cells, filtered through a cell filter mesh, were analyzed using the FACS CANTOII. Live cells were gated based on FSC and SSC properties, and the APC fluorescence intensity of the GFP-positive cell population was measured.

### RT-qPCR Analysis

To coat the bottom of 48-well plates with Notch ligands, 4 ng/µl of rat DLL1-Fc, human DLL4-Fc, or human JAG1-Fc was added to the wells and incubated at room temperature for 2 hours. HEK293T cells, transiently overexpressing pcDNA1 full-length mouse NOTCH1 Myc and pMax-EGFP, were then seeded into the wells. After 24 hours, the cells were washed with PBS, and total RNA was extracted using TRIzol reagent (Thermo). The extracted RNA was reverse-transcribed into cDNA using the PrimeScript RT reagent Kit with gDNA Eraser (Perfect Real Time, Takara). RT-qPCR was performed using SSo Advanced Universal SYBR Green Supermix (Bio-Rad). Relative quantification of gene expression was calculated using the double delta Ct method, with *HES5* expression normalized to the Ct value of *GAPDH*.

### Reporter assay

To coat the bottom of 96-well plates with Notch ligands, 2 ng/µl of mouse DLL1 Myc-His, or mouse DLL4 Myc-His was added to the wells and incubated at 37℃ for 2 hours. HEK293T cells transiently overexpressing pcDNA1 mouse NOTCH1 FL Myc or C^4^-2 site mutants and, TP1-Luc and gWiz-β-Gal were then seeded into the wells. After 24 hours, the cells were washed with PBS and incubated at 22℃ for 10 minutes after addition of Reporter Lysis Buffer (Promega). The lysates were then frozen, thawed, and divided into 96-well white plates and 96-well clear plates.

Luciferase reagent, ONE-Glo EX Buffer (Promega) was added to the white plate, and after incubation at 22℃ for 20 minutes, luminescence was measured using Infinite M200 (TECAN). The β-galactosidase assay buffer [200 mM sodium phosphate buffer pH 7.3, 2 mM MgCl_2_, 100 mM β-mercaptoethanol, 1.33 mg/ml *O*-nitrophenyl β-D-galactopyranoside (Wako)] was added to the transparent plate, incubated at 37℃ for 15 minutes, and the experimental absorbance at 420 nm and the reference absorbance at 630 nm (OD 420/630 nm) were measured using Infinite M200. Relative luciferase units (RLU) were determined by dividing luciferase activity by β-galactosidase activity.

### Statistical Analysis

Multiple analyses were conducted for the results of immunoblotting, flow cytometry, and RT-qPCR. Statistical analysis was performed using R software, and Tukey’s test was applied. Only p-values less than 0.05 were annotated on the graphs.

### Data availability

The glycoproteome data in this study were deposited in jPOSTrepo: an international standard data repository for proteomes (60) (Project ID: JPST003592, PXD060455). NSI-MSn raw data for glycan characterization were deposited into GlycoPost (61), a publicly available data repository (GlycoPost accession number GPST000550).

## Acknowledgments

We thank K. Taki (Division for Medical Research Engineering, Nagoya University Graduate School of Medicine) for supporting the LC-MS/MS analysis, members of the Okajima and Takeuchi laboratories for the critical discussion, and K. Moremen (University of Georgia) and the Repository of Glycoenzyme Expression Constructs (http://glycoenzymes. ccrc.uga.edu/) (the National Institutes of Health Grant P41GM103390 and P01GM107012) for the glycosyltransferase constructs. This work was supported by grants from the Japan Society for the Promotion of Science (JP23KJ1062 to Y.T.; JP19KK0195 and JP19H03176 to H.T.; JP19H03416 and 22H02815 to T.O.), Takeda Science Foundation (to H.T.), and Yamada Science Foundation (to T.O.). The Novo Nordisk Foundation (NNF24OC0088218) and The Danish National Research Foundation (DNRF107) (to Y.N.).

## Competing Interest Statement

The authors have declared no competing interest.

---

**Supporting information table S1.**
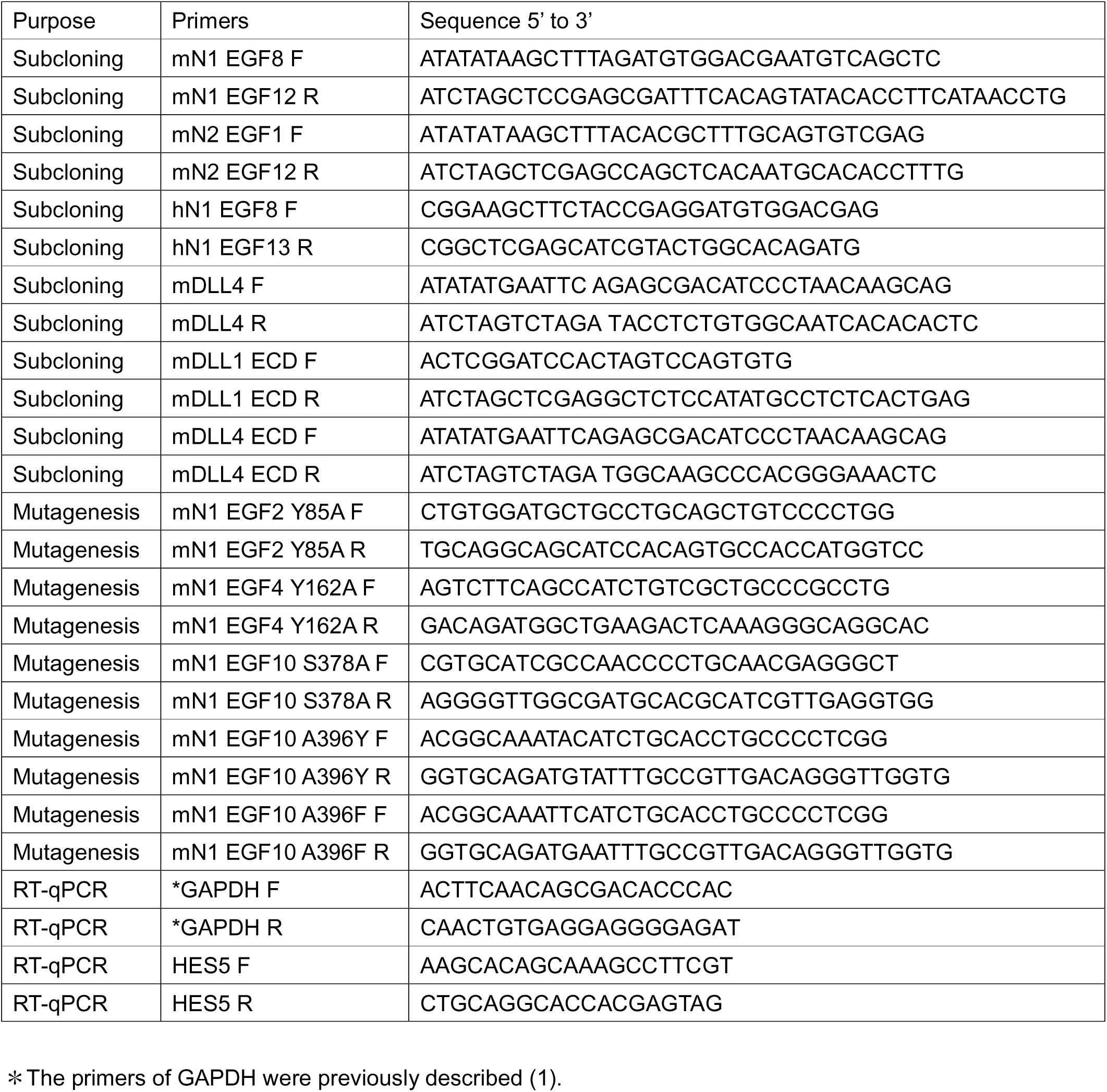
Primers used in this study.

## Supporting information figures

**Fig. S1.** Extracted ion chromatograms (EICs) of mouse NOTCH1 EGF10 expressed in various human cell lines. NOTCH1 fragment proteins were expressed in human cell lines, digested with trypsin, and analyzed by mass spectrometry. EICs of (glyco)peptides, 366-TGLLCHLNDACISNPCNEGSNCDTNPVNGK-395, from mouse NOTCH1 EGF10 are shown. (A) Mouse NOTCH1 EGF6-10 was expressed in a human tongue squamous cell carcinoma cell line, SAS. (B) Mouse NOTCH1 EGF8-12 was expressed in a human pancreatic adenocarcinoma cell line, ASPC1. (C) Mouse NOTCH1 EGF8-12 was expressed in a human acute T-cell leukemia cell line, Jurkat (2). The color coding of EICs is shown in the bottom right. Black: non-*O*-glucosylated peptide; blue: peptide with Glc monosaccharide; yellow: peptide with Xyl-Glc disaccharide; orange: peptide with Xyl-Xyl-Glc trisaccharide; red: peptide with Hex-Glc disaccharide; purple: peptide with Neu5Ac-Hex-Glc trisaccharide.

**Fig. S2.** Glycan composition of *O*-Glc glycans on NOTCH1 EGF10: Effects of *O*-Glc site mutation and xylosyltransferase deficiency/rescue. (A) Mouse NOTCH1 EGF6-10 S378A mutant was expressed in wild-type HEK293T cells, digested with trypsin, and analyzed by mass spectrometry. EICs of mouse NOTCH1 EGF10 tryptic digests, 366-TGLLCHLNDACIANPCNEGSNCDTNPVNGK-395, are shown. (B-C) Wild-type mouse NOTCH1 EGF1-18 was expressed in xylosyltransferases-deficient HEK293T cells, digested with trypsin, and analyzed by mass spectrometry. EICs of mouse NOTCH1 EGF10 tryptic digests, 366-TGLLCHLNDACISNPCNEGSNCDTNPVNGK-395, are shown. B: *GXYLT1/2*-deficient (KO) cells. C: *GXYLT1/2*-deficient cells with *GXYLT1/2* overexpressed (OE). The color coding of EICs is shown in the bottom right. Black: non-*O*-glucosylated peptide; blue: peptide with Glc monosaccharide; yellow: peptide with Xyl-Glc disaccharide; orange: peptide with Xyl-Xyl-Glc trisaccharide; red: peptide with Hex-Glc disaccharide; purple: peptide with Neu5Ac-Hex-Glc trisaccharide.

**Fig. S3.** Full MS spectra of β-eliminated 3′-sialyllactose and the trisaccharide released from NOTCH1 EGF6-10. (A) The standard 3′-sialyllactose was reduced in the presence of sodium borodeuteride (NaBD_4_). Under reduction with NaBD_4_, the standard gained an additional 1 Da compared to the β-eliminated *O*-Glc glycan treated with sodium borohydride (NaBH_4_), as observed in the MS1. (B) Trypsin-digested glycopeptides, 366-TGLLCHLNDACIANPCNEGSNCDTNPVNGK-395, containing the EGF10 *O*-Glc site from NOTCH1 EGF6-10 were subjected to β-elimination with NaBH_4_. Released *O*-Glc glycans were analyzed by mass spectrometry as described in the Materials and methods section.

**Fig. S4.** Glycan composition of *O*-Glc glycans on NOTCH1 EGF10 from *B4GALT1* or *ST3GAL4*-deficient cells rescued by respective gene expression. Wild-type mouse NOTCH1 EGF1-12 was expressed in *B4GALT1*- or *ST3GAL4*-deficient HEK293 cells rescued by overexpression of *B4GALT1* or *ST3GAL4*, respectively. The purified NOTCH1 proteins were digested with trypsin and analyzed by mass spectrometry. EICs of mouse NOTCH1 EGF10 tryptic digests, 366-TGLLCHLNDACISNPCNEGSNCDTNPVNGK-395, are shown. (A) *B4GALT1-*deficient with *B4GALT1* overexpressed. (B) *ST3GAL4-*deficient with *ST3GAL4* overexpressed. The color coding of EICs is shown at the bottom. Black: non-*O*-glucosylated peptide; blue: peptide with Glc monosaccharide; yellow: peptide with Xyl-Glc disaccharide; orange: peptide with Xyl-Xyl-Glc trisaccharide; red: peptide with Gal-Glc disaccharide; purple: peptide with Neu5Ac-Gal-Glc trisaccharide.

**Fig. S5.** Glycan composition of *O*-Glc glycans on NOTCH1 C^4^-2 site-mutants. (A-E) Wild-type (WT) or four different C^4^-2 mutants of mouse NOTCH1 EGF1-12 were expressed in wild-type HEK293T cells. The purified NOTCH1 proteins were digested with trypsin and analyzed by mass spectrometry. EICs of tryptic digests, 57-CQDSNPCLSTPCK-69, 137-SCQQADPCASNPCANGGQCLPFESSYICR-165, or 366-TGLLCHLNDACISNPCNEGSNCDTNPVNGK-395, from EGF2, EGF4, or EGF10, respectively, were shown. A: WT. B: A396Y mutant (C^4^-2 of EGF10). C: A396F mutant (C^4^-2 of EGF10). D: Y85A mutant (C^4^-2 of EGF2). E: Y162A mutant (C^4^-2 of EGF4) (See also glycan composition data at EGF10 from WT or A396Y and A396F in Fig. 1 or Fig. 4, respectively.). The color coding of EICs is shown in the bottom right. Black: non-*O*-glucosylated peptide; blue: peptide with Glc monosaccharide; yellow: peptide with Xyl-Glc disaccharide; orange: peptide with Xyl-Xyl-Glc trisaccharide; red: peptide with Gal-Glc disaccharide; purple: peptide with Neu5Ac-Gal-Glc trisaccharide.

**Fig. S6.** Impact of C^4^-2 site mutations on NOTCH1 protein expression. Mutations at two amino acids before the fourth cysteine were introduced to mouse NOTCH1 EGF 1-12 or full-length mouse NOTCH1. (A) Western blotting analysis of secreted mouse NOTCH1 EGF1-12. Mouse anti-Myc antibody was used as a primary antibody. HRP-conjugated goat anti-mouse IgG-Fc was used as a secondary antibody. *Top*: mouse NOTCH1 EGF1-12; *Bottom*: IgG-Fc co-transfected as a transfection control for normalization. (B) Flow cytometry analysis of full-length mouse NOTCH1 cell surface expression. APC-conjugated anti-mouse NOTCH1 antibody was used. Viable cells were gated using SSC and FSC parameters, and transfected cells were further selected based on co-expressed GFP fluorescence. Red: WT; blue: A396Y; yellow: A396F.

**Fig. S7.** Analysis of glycan composition and ligand binding properties in mouse NOTCH1 EGF8-12 variants. (A) Glycan composition of mouse NOTCH1 EGF8-12, produced in *POGLUT1*-overexpressing wild-type HEK293T cells, containing both galactose- and xylose-elongated *O*-Glc glycans. (B) Glycan composition of mouse NOTCH1 EGF8-12, produced in *POGLUT1*- and *GXYLT1*-overexpressing *B4GALT1* KO cells, lacking galactose elongated *O*-Glc glycans. (C) Glycan composition of mouse NOTCH1 EGF8-12, produced in *POGLUT1*- and *B4GALT1*-overexpressing *GXLYT1/2* KO cells, lacking xylose-elongated *O*-Glc glycans. The color coding of EICs is shown at the bottom. Black: non-*O*-glucosylated peptide; blue: peptide with Glc monosaccharide; yellow: peptide with Xyl-Glc disaccharide; orange: peptide with Xyl-Xyl-Glc trisaccharide; red: peptide with Gal-Glc disaccharide; purple: peptide with Neu5Ac-Gal-Glc trisaccharide. (D-F) Bar graphs showing NOTCH1 (5 ng/μl) binding to each ligand. D: DLL1-expressing cells. E: DLL4-expressing cells. F: JAG1-expressing cells. Circles: mouse NOTCH1 EGF8-12 whose glycan composition is shown in (A); squares: mouse NOTCH1 EGF8-12 whose glycan composition is shown in (B); triangles: mouse NOTCH1 EGF8-12 whose glycan composition is shown in (C). Statistical analysis was performed using one-way ANOVA followed by Tukey’s test. Error bars represent SEM.

**Fig. S8.** Binding of Notch ligands to NOTCH1-expressing cells. Wild-type (WT) or C^4^-2 site-mutated full-length mouse NOTCH1 were expressed in HEK293T cells. The cells were incubated with recombinant Notch ligand-Fc proteins. Concentrations of Notch ligands are indicated. (A) DLL1 binding (N = 3). (B) DLL4 binding (N = 3). (C) JAG1 binding (N = 3). Black line with a white circle: full-length mouse NOTCH1 WT; black line with a black square: full-length mouse NOTCH1 A396Y mutant; black line with a black triangle: full-length mouse NOTCH1 A396F mutant. Relative mean fluorescence intensity is shown on the y-axis. The intensity of WT is set as 1.0. (D-F) Bar graphs showing binding of DLL1 (D), DLL4 (E), or JAG1 (F) (each at 4 ng/μl) to NOTCH1-expressing cells. Statistical analysis was performed using one-way ANOVA followed by Tukey’s test. Error bars represent SEM.

