## Supplementary figures and images for "Differential *O*-Glc elongation on the specific EGF repeat within a ligand binding domain regulates NOTCH1 signaling"

### Fig S1

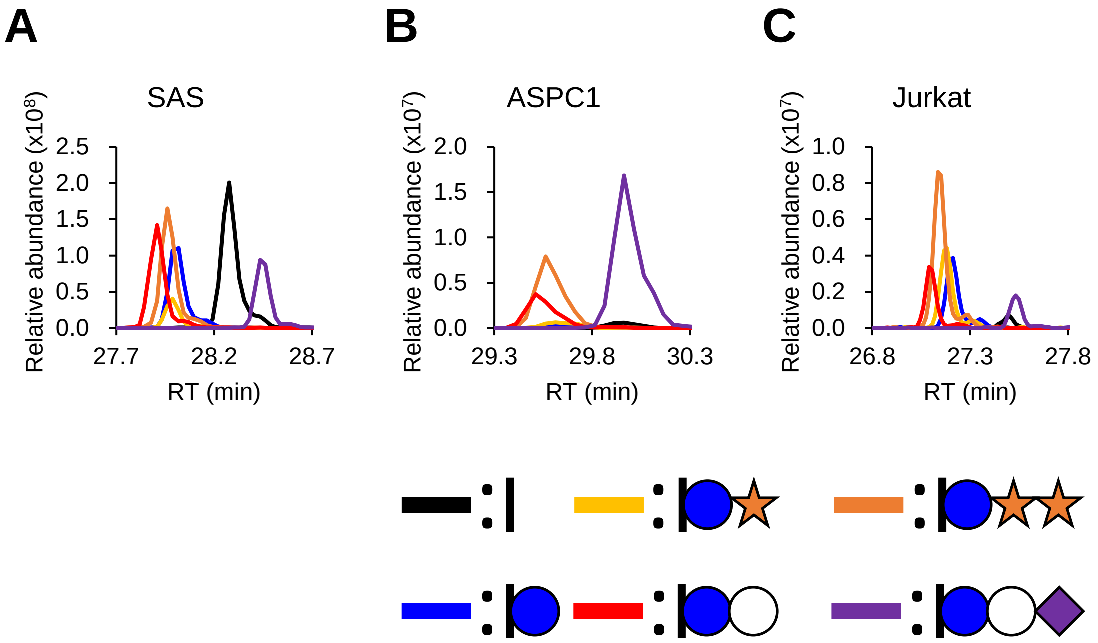

### Fig S2

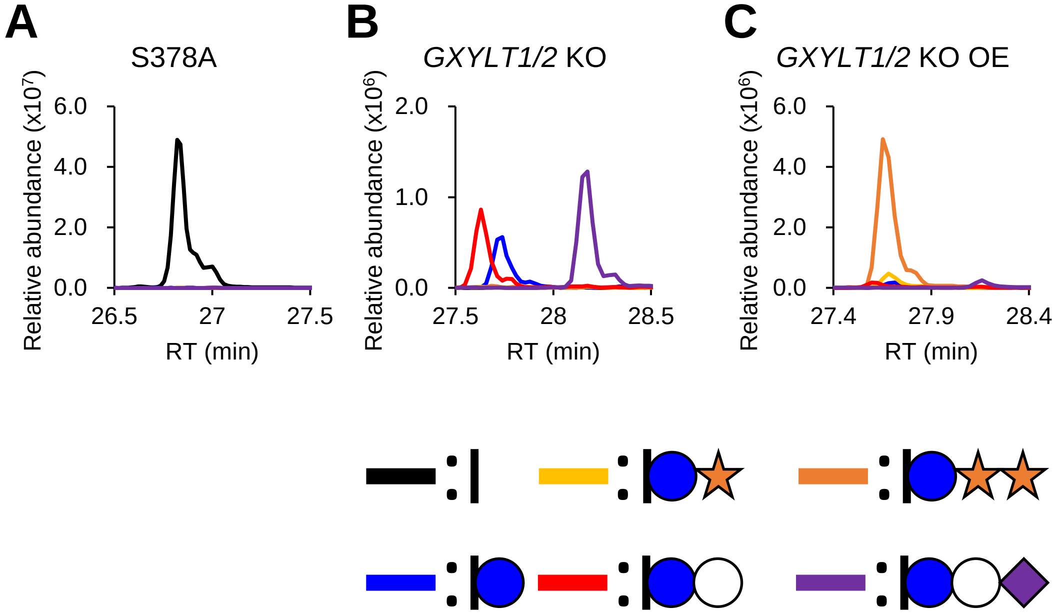

### Fig S3

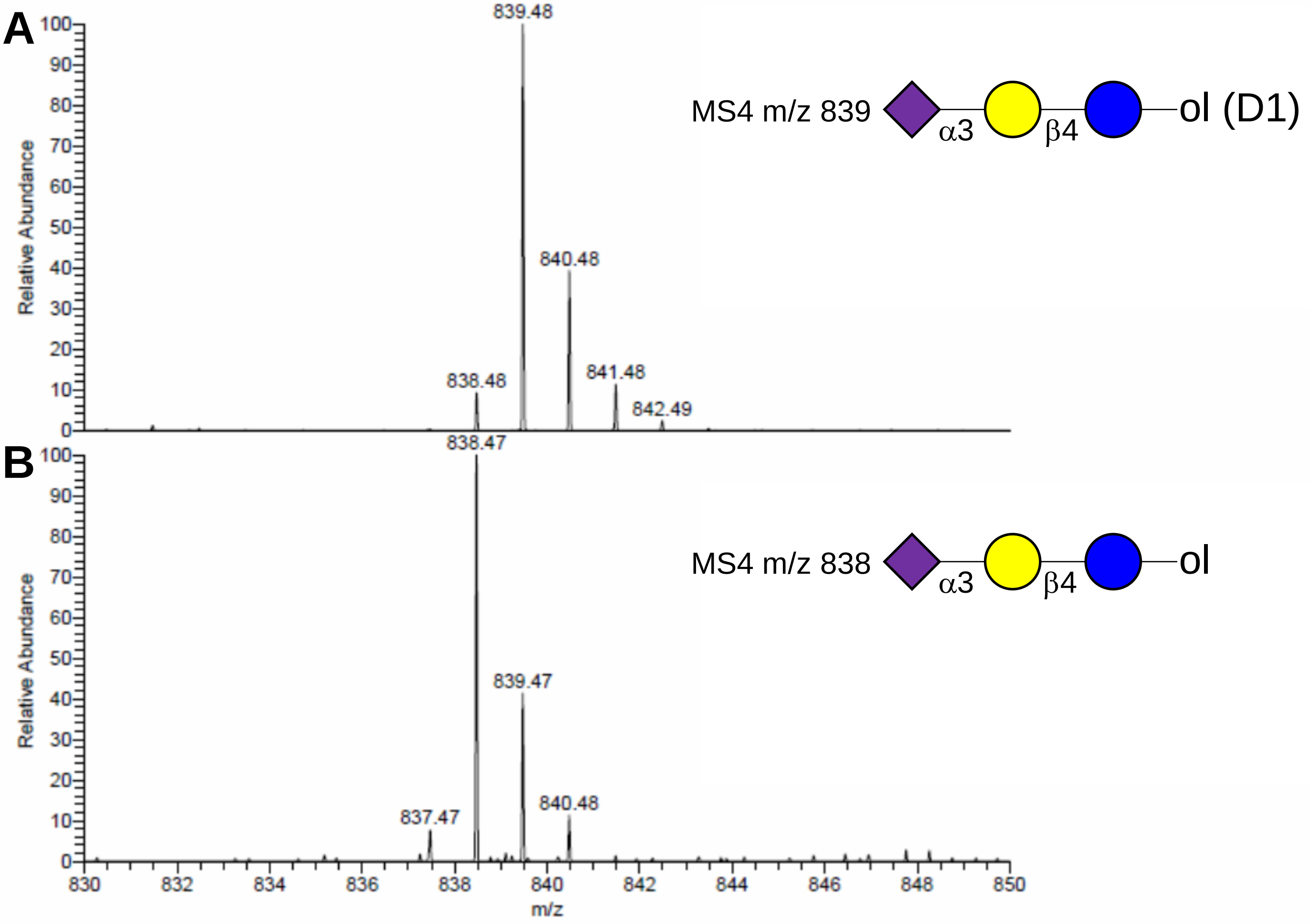

### Fig S4

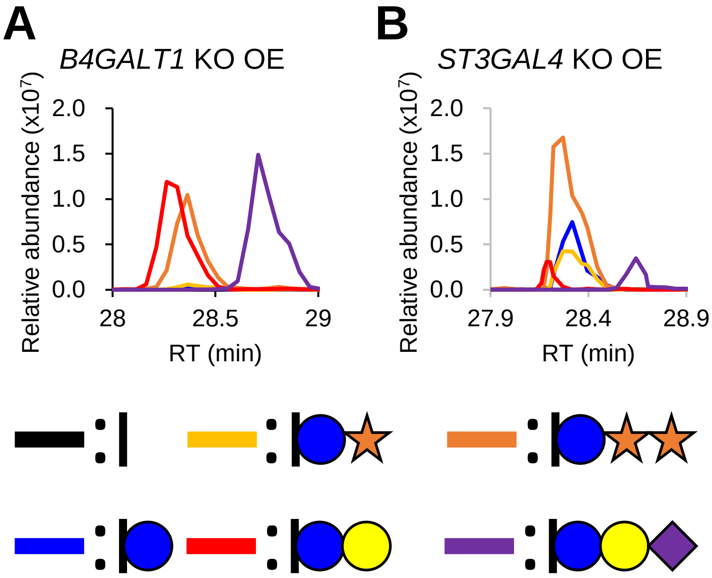

### Fig S5

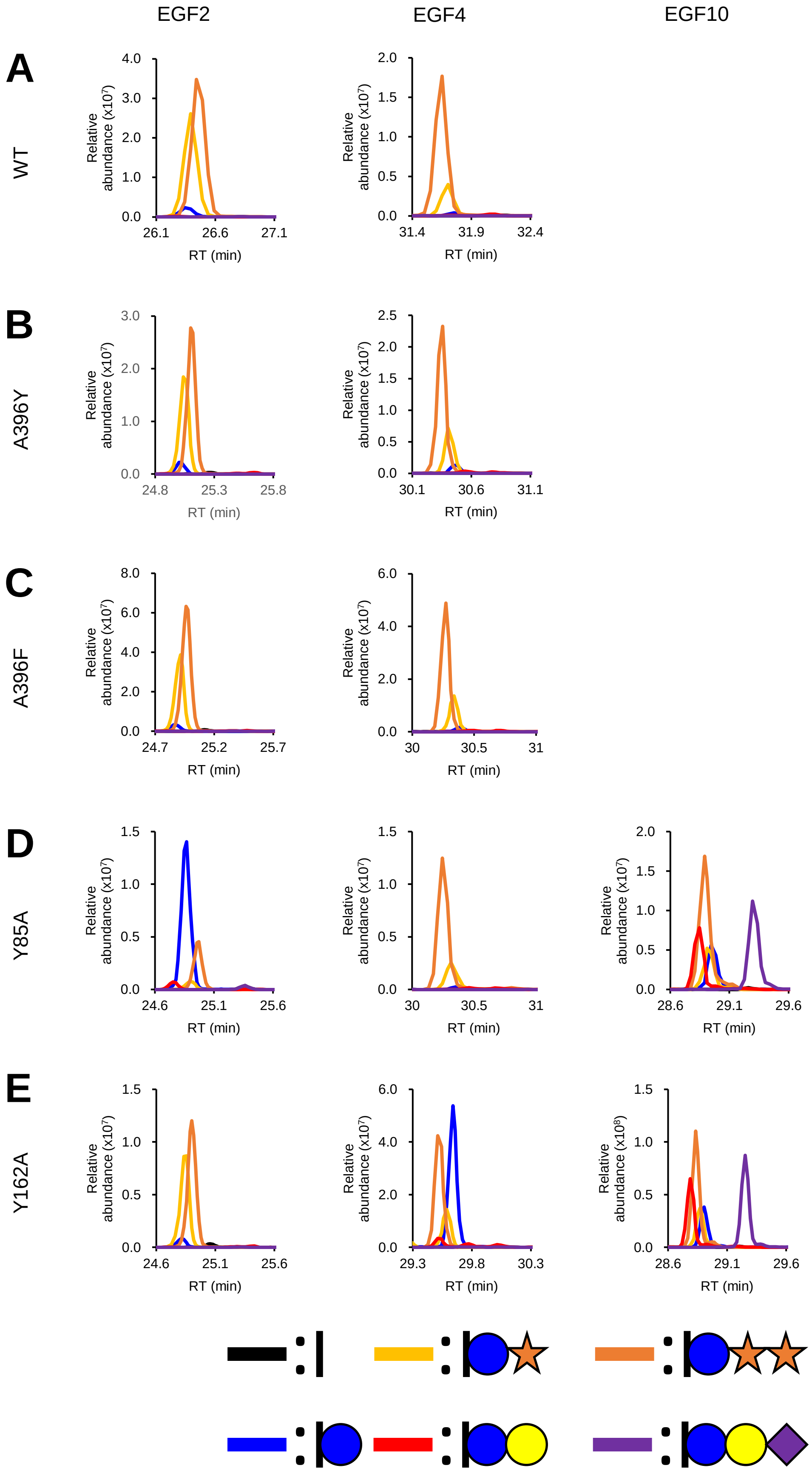

### Fig S6

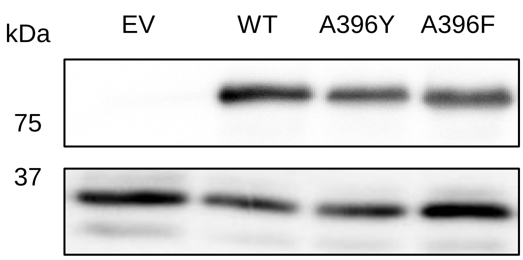

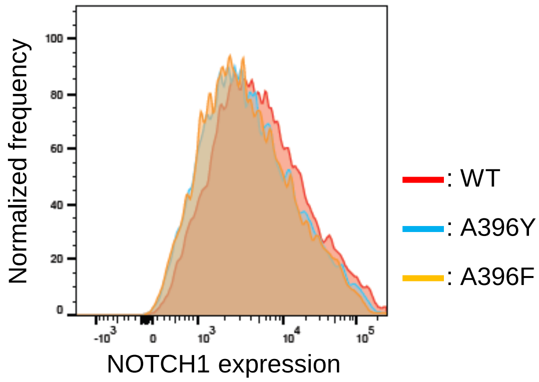

### Fig S7

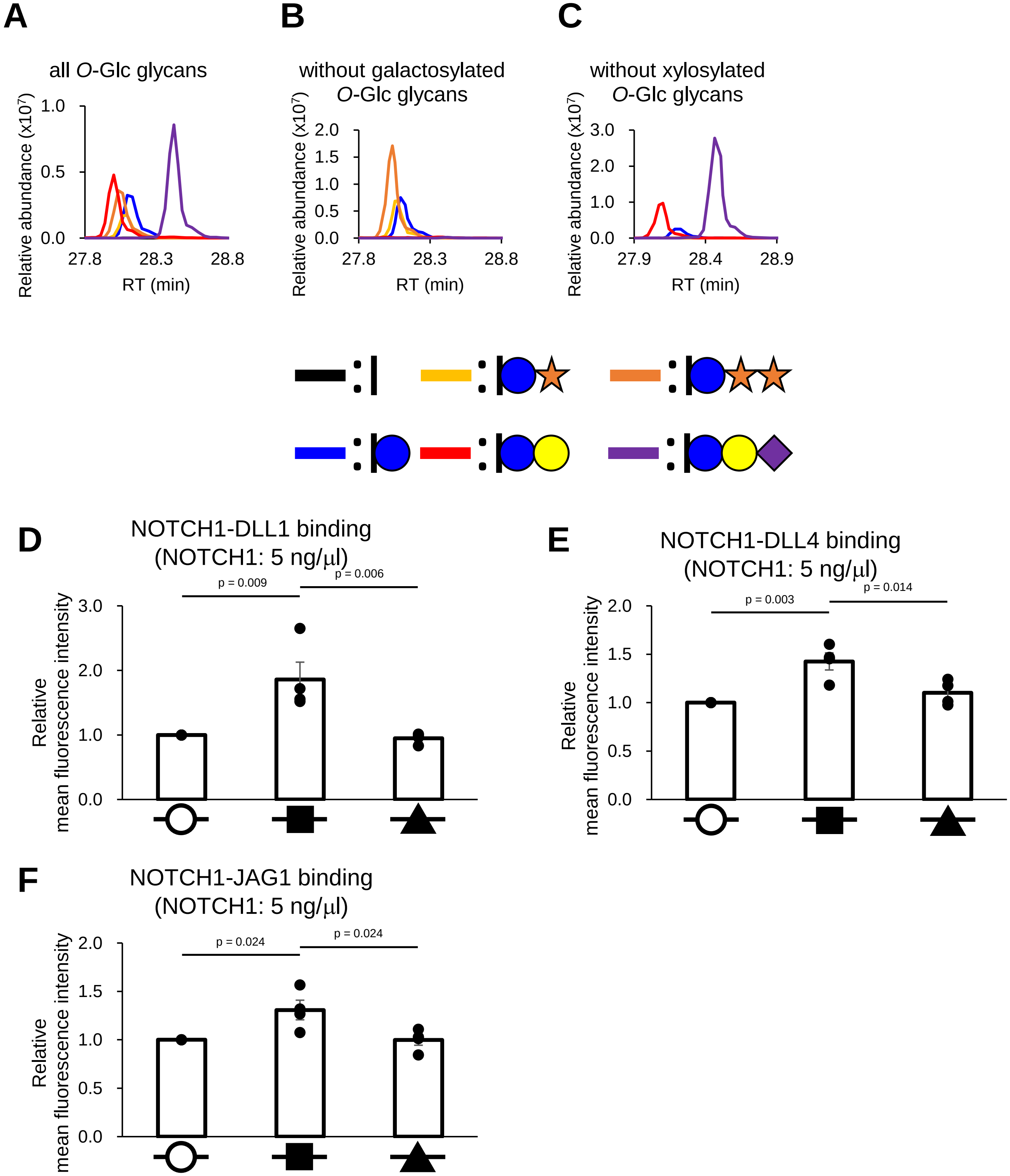

### Fig S8

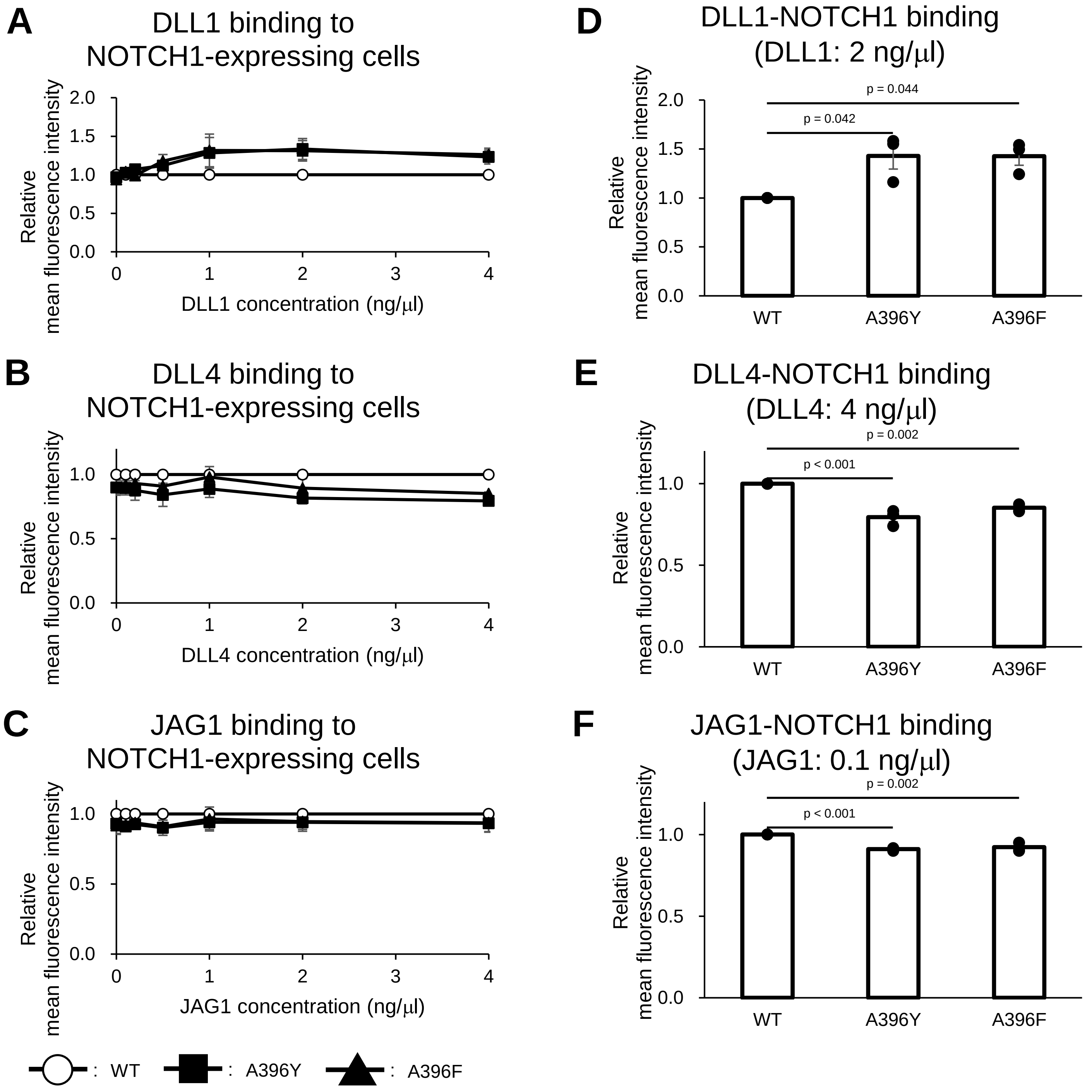
